# Cholesterol Dependence of the Conformational Changes in Metabotropic Glutamate Receptor 1

**DOI:** 10.1101/2024.04.17.589854

**Authors:** Ugochi H. Isu, Shadi A. Badiee, Adithya Polasa, Seyed H. Tabari, Mortaza Derakhshani-Molayousefi, Mahmoud Moradi

## Abstract

Metabotropic glutamate receptors (mGluRs) are class C G protein-coupled receptors that function as obligate dimers in regulating neurotransmission and synaptic plasticity in the central nervous system. The mGluR1 subtype has been shown to be modulated by the membrane lipid environment, particularly cholesterol, though the molecular mechanisms remain elusive. In this study, we employed all-atom molecular dynamics simulations to investigate the effects of cholesterol on the conformational dynamics of the mGluR1 seven-transmembrane (7TM) domain in an inactive state model. Simulations were performed with three different cholesterol concentrations (0%, 10%, and 25%) in a palmitoyl-oleoyl phosphatidylcholine (POPC) lipid bilayer system. Our results demonstrate that cholesterol induces conformational changes in the mGluR1 dimer more significantly than in the individual protomers. Notably, cholesterol modulates the dynamics and conformations of the TM1 and TM2 helices at the dimer interface. Interestingly, an intermediate cholesterol concentration of 10% elicits more pronounced conformational changes compared to both cholesterol-depleted (0%) and cholesterol-enriched (25%) systems. Specific electrostatic interaction unique to the 10% cholesterol system further corroborate these conformational differences. Given the high sequence conservation of the 7TM domains across mGluR subtypes, the cholesterol-dependent effects observed in mGluR1 are likely applicable to other members of this receptor family. Our findings provide atomistic insights into how cholesterol modulates the conformational landscape of mGluRs, which could impact their function and signaling mechanisms.

## Introduction

The intricate interplay between membrane lipids and integral membrane proteins plays a crucial role in regulating various cellular processes, from signal transduction to membrane trafficking.^1,2^ Among the diverse array of membrane lipids, cholesterol stands out due to its unique physicochemical properties and functional versatility.^3^ Cholesterol not only modulates the biophysical properties of cell membranes, such as fluidity and permeability, but also engages in specific interactions with membrane proteins, influencing their structure, stability, and activity.^4–7^ Deciphering the complex relationship between cholesterol and membrane proteins is therefore essential for understanding the molecular mechanisms underlying cellular signaling and function.

G protein-coupled receptors (GPCRs) constitute a prominent class of membrane proteins that mediate a wide range of physiological responses to extracellular stimuli.^8,9^ Within the GPCR superfamily, metabotropic glutamate receptors (mGluRs) are key regulators of synaptic transmission and neuronal excitability in the central nervous system.^10–13^ As members of the Class C GPCR family, mGluRs exhibit a unique structural architecture, comprising a large extracellular Venus Flytrap domain (VFT) responsible for ligand binding, coupled to a seven-transmembrane helical bundle (7TM) that transduces signaling events across the membrane.^4,14–17^ The activation of mGluRs triggers a cascade of intracellular signaling events,^18–20^ modulating synaptic plasticity and neuronal function.^21–23^

Given the vital role of mGluRs in synaptic transmission and neuronal function, elucidating the factors that regulate their structure and function is of utmost importance. Recent studies have shed light on the significance of membrane lipids, particularly cholesterol, in modulating mGluR structure and function. ^4,16,24–26^ For instance, cholesterol depletion has been shown to impair the function of mGluR1 in hippocampal neurons, leading to reduced calcium signaling and altered synaptic plasticity.^24,27,28^ Moreover, cholesterol has been implicated in the localization and trafficking of mGluRs, with cholesterol-rich lipid rafts serving as platforms for mGluR signaling complexes. ^16,29–32^ These findings suggest a potential role for lipid-protein interactions in fine tuning mGluR signaling.

Despite these insights, the molecular mechanisms underlying cholesterol-dependent modulation of mGluRs, particularly mGluR1, remain poorly understood. Investigating the impact of cholesterol on the conformational dynamics of mGluR1 is necessary for unraveling the molecular basis of mGluR signaling and may have implications for the development of novel therapeutic strategies targeting mGluRs in neurological disorders.

In this study, we employ molecular dynamics (MD) simulations to investigate the relationship between cholesterol dependence and mGluR1 conformational changes, aiming to gain insights into this dynamic process. MD simulations have emerged as a powerful tool for probing protein-lipid interactions at the atomic level, providing invaluable insights into the dynamic behavior of membrane proteins. ^33–36^ Our findings suggest that cholesterol could stabilize specific conformational states of mGluR1, potentially impacting its activation and signaling properties. We show that cholesterol binding to specific regions on mGluR1 could modify the receptor’s conformational landscape, possibly favoring conformations conducive to G protein coupling and subsequent downstream signaling events. The insights gained from this study not only advance our understanding of the complex interplay between cholesterol and mGluR1 but also have broader implications for the role of lipid-protein interactions in regulating GPCR signaling. By shedding light on the molecular mechanisms behind cholesterol-dependent modulation of mGluR1, our findings could contribute to a better understanding and offer potential avenues for developing therapeutic strategies targeting mGluRs in neurological disorders like schizophrenia and Parkinson’s disease, where abnormal mGluR signaling is implicated.^37–44^

## Methods

We conducted all-atom MD simulations to investigate the influence of cholesterol on the conformational changes of mGluR1 using a homogeneous lipid bilayer consisting of pure 1-palmitoyl-2-oleoyl-sn-glycero-3-phosphocholine (POPC) and heterogeneous lipid bilayers consisting of 10% cholesterol and 90% POPC), and 25% cholesterol and 75% POPC. We constructed two sets of three mGluR1 systems, excluding the bound thermostabilized apocytochrome *b*_5_62 (BRIL) protein from the crystal structure in both sets. The first set incorporated the six molecules of cholesterol from the crystal structure into the 10% and 25% systems, whereas the second set had these initial cholesterol molecules removed. Simulations were set up using the CHARMM-GUI platform,^45–47^ utilizing the crystal structure of the human class C G protein-coupled mGluR1 in complex with a negative allosteric modulator (PDB entry: 4OR2).^48^ Cholesterol molecules present in the crystal structure were removed for the 0% system in both sets, while the protein was placed in lipids, solvated in a box of TIP3P waters, and 0.15M NaCl using CHARMM-GUI.^45–47^ The box size for the systems was approximately *≈* 150 Å *×* 150 Å *×* 130 Å*and ≈* 84 Å *×* 84 Å *×* 112 Å for both systems, with a total of 152713, 196337, and 192820 atoms for 0%, 10%, and 25%, respectively in set1, and 71848, 73930, and 73260 atoms for 0%, 10%, and 25% respectively in set2, and all systems were simulated in apo conditions (that is without the bound negative allosteric modulator from the crystal structure).^48^ The final systems contained the following lipid compositions: For Set 1, there were 598 lipids, with 298 lipids in the upper leaflet and 300 lipids in the lower leaflet for 0% cholesterol. For 10% cholesterol, there were 630 lipids, including 31 cholesterol and 279 POPC lipids in the upper leaflet, and 32 cholesterol and 288 POPC lipids in the lower leaflet respectively. For 25% cholesterol, there were 668 lipids, consisting of 83 cholesterol and 249 POPC lipids in the upper leaflet, and 84 cholesterol and 252 POPC lipids in the lower leaflet respectively. For Set 2, there were 232 lipids, with 115 lipids in the upper leaflet and 117 lipids in the lower leaflet for 0% cholesterol. For 10% cholesterol, there were 250 lipids, comprising 12 cholesterol and 108 POPC lipids in the upper leaflet, and 13 cholesterol and 117 POPC lipids in the lower leaflet. For 25% cholesterol, there were 250 lipids, including 32 cholesterol and 96 POPC lipids in the upper leaflet, and 33 cholesterol and 99 POPC lipids in the lower leaflet.

All systems were simulated with NAMD 2.10/2.13^49^ and the CHARMM36m all-atom additive force field^50^.^51^ Initially each system was energy-minimized for 10,000 steps using the conjugate gradient algorithm.^52^ Then, we relaxed the systems by applying restraints in a stepwise manner (for a total of *∼*1ns) using the standard CHARMM-GUI equilibration protocol.^53^ Production runs were carried out for 960ns each and 1 *µ*s each for both sets respectively. The initial relaxation was performed in an NVT ensemble, while all production runs were performed in an NPT ensemble. Simulations were conducted at 310 K using a Langevin integrator with a damping coefficient of *γ* = 0.5 *ps^−^*^1^. The pressure was maintained at 1atm using the Nośe-Hoover Langevin piston method.^52,54^ The smoothed cutoff distance for non bonded interactions was set to 10-12 Å, and long-range electrostatic interactions were computed with the particle mesh Ewald (PME) method.^55^

The TM helices and other subdomains were defined as follows: TM1 (residue 592 - 616), TM2 (629 - 647), TM3 (654- 683), TM4 (703 - 727), TM5 (753-772), TM6 (784 - 809), TM7 (812 - 840), ICL1 region (617 -628), ICL2 loop (681-702), ICL3 (773-783), ECL1(648-653), ECL2(731-745), ECL3(810-811).^48^ The root mean square deviation (RMSD) trajectory tool of VMD^56–58^ was used to calculate the RMSD, with C*α* atoms considered for these calculations. Root mean square fluctuation (RMSF) of individual residues was calculated using C*α* atoms, and the VMD timeline plugin was employed to identify salt bridges. Lipid-protein interactions were characterized by counting the number of lipid molecules within 4 Å of the protein or specific protein domains at every frame. Principal component analysis (PCA) was conducted using PRODY software,^59^ considering only C*α* atoms. Hydrogen bond analysis was performed using the VMD HBond plugin, with a cutoff distance of 3.5 Å and an angle of 30*^◦^*. Dynamical network analysis was done using VMD^60^ and Carma.^61^

## Results and Discussion

### Cholesterol Influences the Conformational Changes of the Internal Protein and Acts Less Significantly on Individual Protomers

To provide mechanistic insights into cholesterol sensitivity for the mGluR1 receptor, PCA was employed to identify the most significant differences in conformational dynamics between the three studied systems: mGluR1 in the presence of 0%, 10%, and 25% cholesterol. The projections of the MD trajectories onto the first two principal components (PC1 and PC2) reveal distinct behaviors of individual protomers and the entire mGluR1 protein (Fig. 2A-C). The PCA results demonstrate that the most pronounced conformational changes captured by PC1, exhibit significant variations between the cholesterol-containing systems (10% and 25%CHOL) and the cholesterol-free system (0% CHOL). Notably, the system without cholesterol displays reduced motion along PC1, indicating a more restricted conformational space (Fig. 2C). The differences in conformational dynamics are more evident when considering the entire mGluR1 protein compared to individual protomers. This observation suggests that cholesterol exerts a more pronounced influence on the interprotomer conformational dynamics rather than the dynamics of single protomers. The presence of cholesterol appears to modulate the collective motions and coordinated conformational changes of the mGluR1 dimeric assembly. We also observe from the PCA results that the system with 10% cholesterol exhibits higher conformational variability compared to the system with 0% and 25% cholesterol (Fig. 2C) when considering the whole protein structural conformation. This finding implies that increasing cholesterol concentration may induce a more ordered and less dynamic structure of mGluR1. It is plausible that higher cholesterol levels promote stronger interactions between the receptor and the lipid bilayer, constraining the conformational flexibility of the protein.^62–64^

**Fig. 1.**
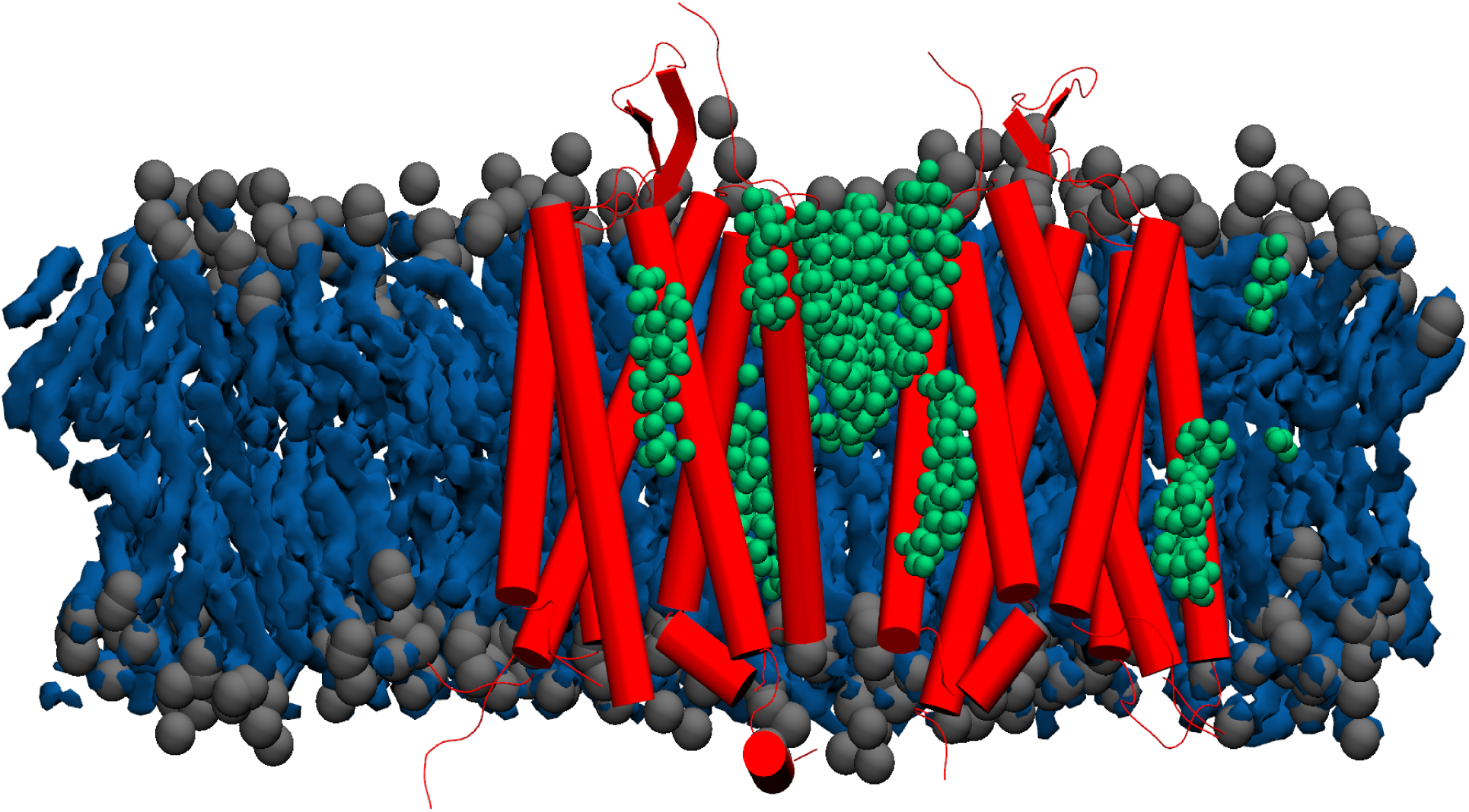
A cartoon representation of mGluR1 (PDB: 4OR2) illustrates the interaction sites of cholesterol within the protein embedded in a lipid membrane (blue with gray headgroups). Cholesterol molecules (green) are shown interacting between the monomers of mGluR1 (highlighted in red) and within the grooves of the transmembrane helices.

**Fig. 2.**
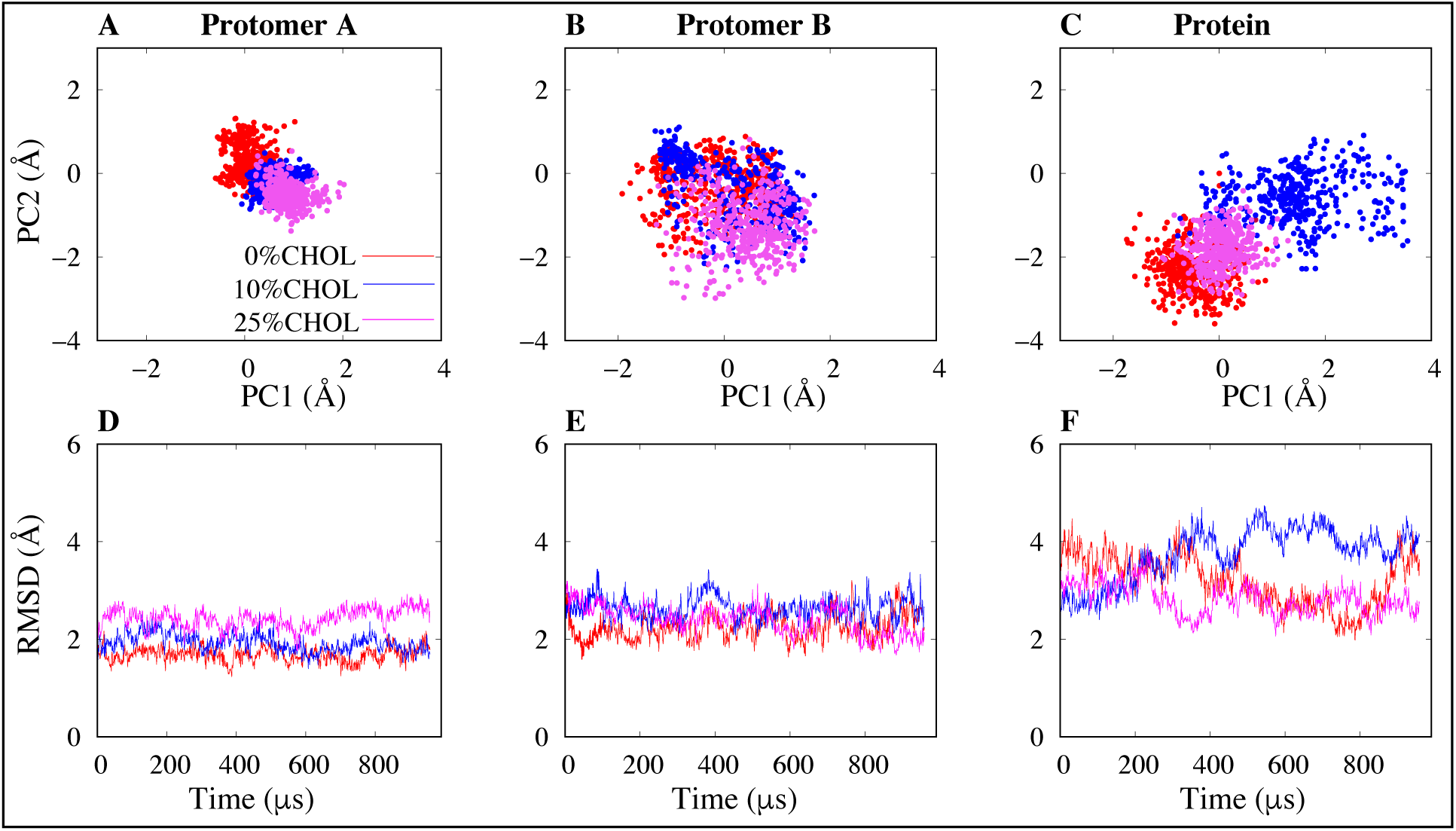
Projections of the principal components (PC’s) 1 and 2 [A-C], and the root mean square deviation analysis [D-F] of mGluR1 in the presence and absence of cholesterol, in the first simulation set.

To assess the structural stability of mGluR1 during the simulations, we calculated the RMSD of the protein backbone (Fig. 2D-F) for the final 1000 frames. Our results show that the cholesterol-free system (0% CHOL) exhibits lower RMSD values compared to the systems with 10% and 25% cholesterol in protomer A and B, however the highest conformational change is observed when the entire protein structure is considered and more in the 10% system.

The combined analysis of PCA and RMSD results (Fig. 2) reveals that cholesterol significantly influences the global conformational stability and dynamics of mGluR1. The presence of cholesterol induces distinct conformational changes in the overall protein structure, while its impact on individual protomers appears to be less pronounced. These findings highlight the importance of considering the lipid environment, particularly cholesterol, when studying the functional dynamics of mGluR1 and other membrane proteins. ^62^ The observed cholesterol-dependent modulation of mGluR1 conformational dynamics could have implications for understanding the receptor’s function and its potential as a therapeutic target. The interaction between cholesterol and mGluR1 may influence the receptor’s ability to bind ligands, undergo conformational changes, and initiate downstream signaling cascades.^65,66^

### Low Cholesterol Concentration (10% CHOL) Induces a Higher Conformational Change

The previous analysis suggested that 10% cholesterol exhibits higher conformational variability compared to the system with 0% and 25% cholesterol (Fig. 2C & F). To confirm this, we investigated the influence of cholesterol on the inter-protomer distance and angle of mGluR1. From our result, the inter-protomer distance showed the lowest distance in the cholesterol-free system (0% CHOL), with the system with higher cholesterol (25% CHOL) displaying a relatively similar behavior to 0% CHOL. The inter-protomer distance of 0% CHOL maintains a distance of approximately 38-39 Å throughout the simulation, while 25% fluctuates from 39-40 Å (Fig. 3A). However, the system containing 10% cholesterol displays a distinct behavior compared to the other two systems. The inter-protomer distance in the 10% cholesterol system undergoes significant fluctuations, reaching a minimum of around 38 Å and a maximum of approximately 42 Å (Fig. 3A). This observation indicates that an intermediate cholesterol concentration induces a more dynamic and flexible arrangement of the mGluR1 protomers, allowing for a wider range of conformational states. The similar behavior of the cholesterol-free and 25% cholesterol systems suggests that the mGluR1 dimer maintains a stable and consistent conformation in the absence of cholesterol and at high cholesterol concentrations. This finding implies that the impact of cholesterol on the inter-protomer distance is not linear and may exhibit a concentration-dependent effect.

**Fig. 3.**
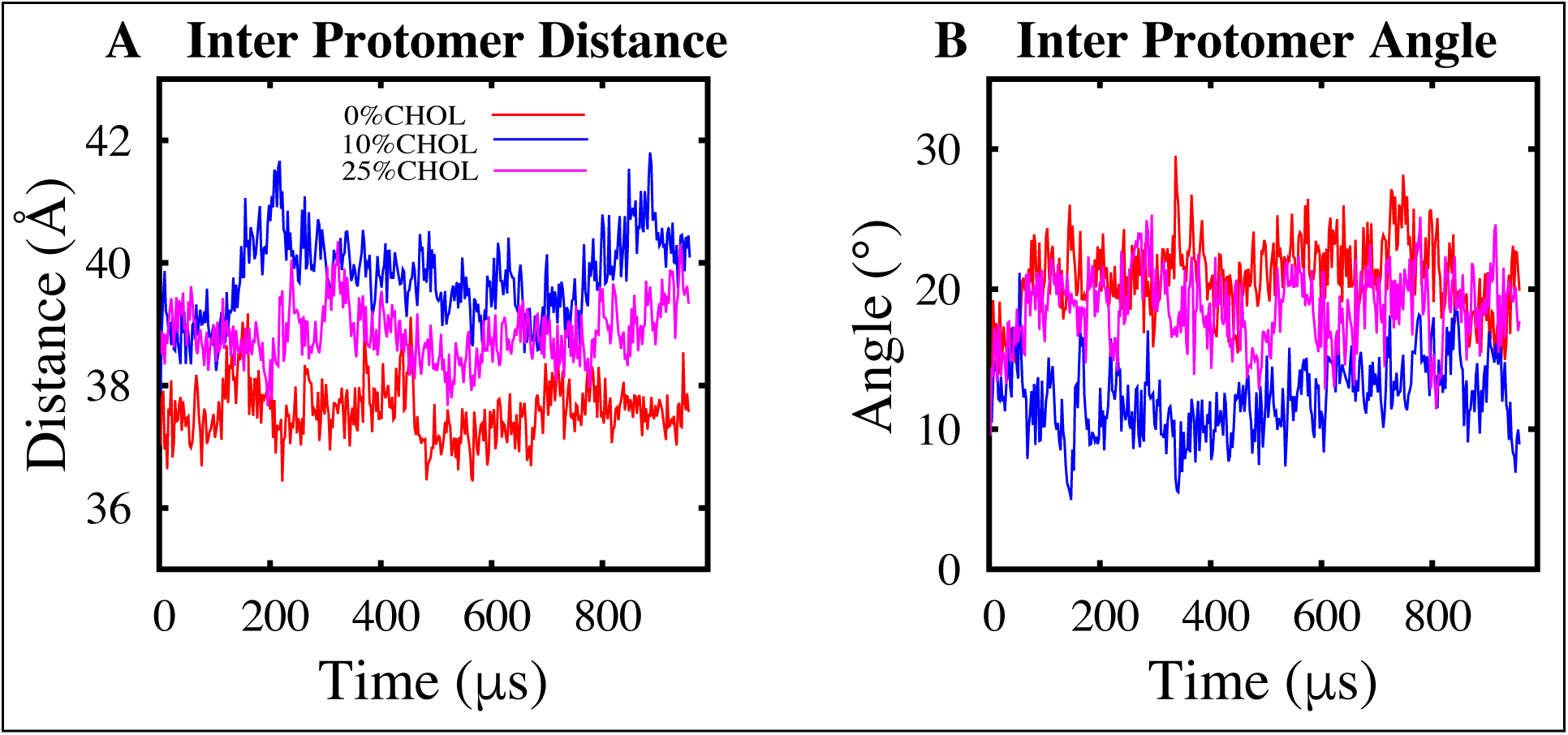
Time series representation of the inter protomer distance (left panel) and inter protomer angle (right panel) in simulation set 1.

The analysis of the inter-protomer angle (Fig. 3B) further supports the distinct behavior of the 10% cholesterol system. While the cholesterol-free and 25% cholesterol systems show higher inter-protomer angles throughout the simulation, the 10% cholesterol system showed the lowest angle, ranging from approximately 20*^◦^* to 5*^◦^* (Fig. 3B). This indicates that an intermediate cholesterol concentration promotes a more dynamic and flexible relative orientation between the mGluR1 protomers.

In addition, the water density maps show the average water occupancy within the simulation system during the initial and final 200 nanoseconds of the trajectories. Within the 0% CHOL system, the water density maps reveal a relatively consistent distribution of water molecules surrounding the mGluR1 protein throughout the simulation. The protein exhibits a uniform pattern of water occupancy in both the initial and final frames of the trajectory, but with a slightly higher water density in the final frames.(Fig. 4). However, when the cholesterol concentration is increased to 10%, a notable change in the water density distribution is observed. During the initial 200 nanoseconds, we observe a significant water occupancy, but in the final 200 nanoseconds, a distinct shift in the water density pattern is observed. The water occupancy within specific regions of the protein seems to decrease, suggesting a reorganization or exclusion of water molecules from these areas (Fig. 4). This observation contrasts with the more consistent water density patterns observed in the systems with 0% and 25% cholesterol. The system containing 25% cholesterol displays a relatively stable water density distribution throughout the entire simulation (Fig. 4), indicating a more persistent and uniform environment surrounding the mGluR1 receptor. To summarize, comparing the initial frames of the first 200 nanoseconds to the final frames, we observed a slight increase in water occupancy at 0%, a decrease at 10%, and a relative uniformity at 25%. These localized alterations in water occupancy around the receptor could be indicative of specific structural rearrangements or the formation of cholesterol-mediated interactions that influence the protein’s conformational landscape.

**Fig. 4.**
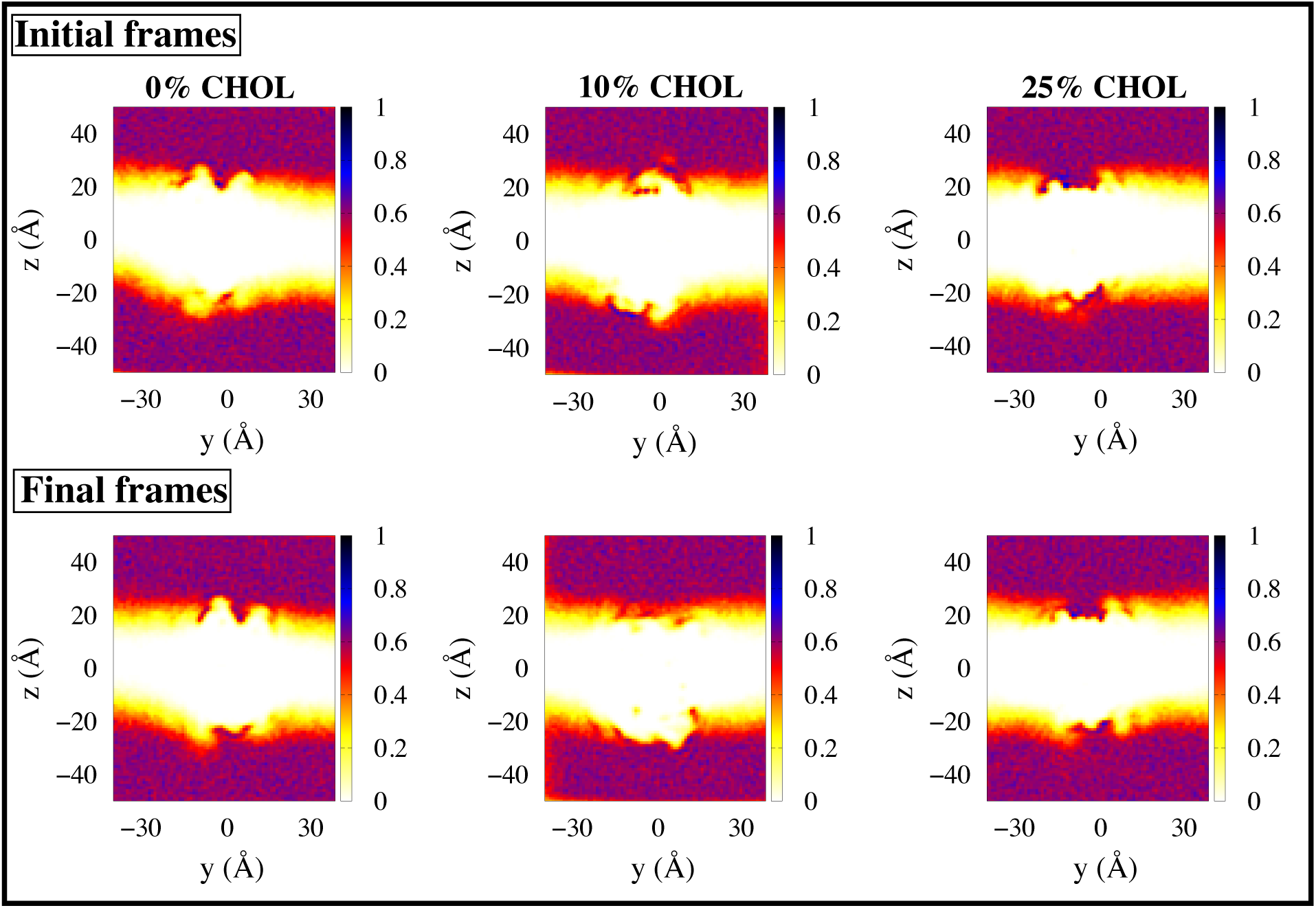
Water density maps illustrate the initial frame (first 200 ns) and final frames (last 200 ns) for Set 1 across 0%, 10%, and 25% cholesterol concentrations. Variations in correlation are represented by a color gradient, ranging from 0 (pale yellow) to 1 (dark blue).

The analysis of the salt bridge interaction distance between Glu 728 and Arg 661 in Protomer A and B (Fig. 5A & B) provides additional insights into the cholesterol-mediated effects on the mGluR1 receptor. Our results reveal that the Glu 728-Arg 661 distance is most stable in the presence of 10% cholesterol in Protomer A. In this system, the salt bridge interaction is not initially present in the crystal structure, but forms at around 200 nanoseconds and is then maintained for the duration of the simulation (Fig. 5A), indicating the development and persistence of this structural feature. In contrast, the cholesterol-free system (0% CHOL) and the 25% cholesterol system exhibit more fluctuations in the Glu 728 - Arg 661 distance within Protomer A (Fig. 5A). This suggests that the absence of cholesterol and the presence of a high cholesterol concentration may lead to a less stable interaction between these critical amino acid residues in this particular protomer. However, the observations in Protomer B reveal a different pattern. In the 0% cholesterol system, a less stable interaction between Glu 728 and Arg 661 is observed for about 300 nanoseconds. Notably, this salt bridge interaction is not present in any of the cholesterol-rich systems (10% and 25% CHOL) within Protomer B (Fig. 5B). These findings suggest that an intermediate cholesterol concentration of 10% may contribute to the overall structural integrity and functional organization of the mGluR1 receptor, potentially through the stabilization of key interresidue interactions (Tab. S1), such as the Glu 728-Arg 661 salt bridge.

**Fig. 5.**
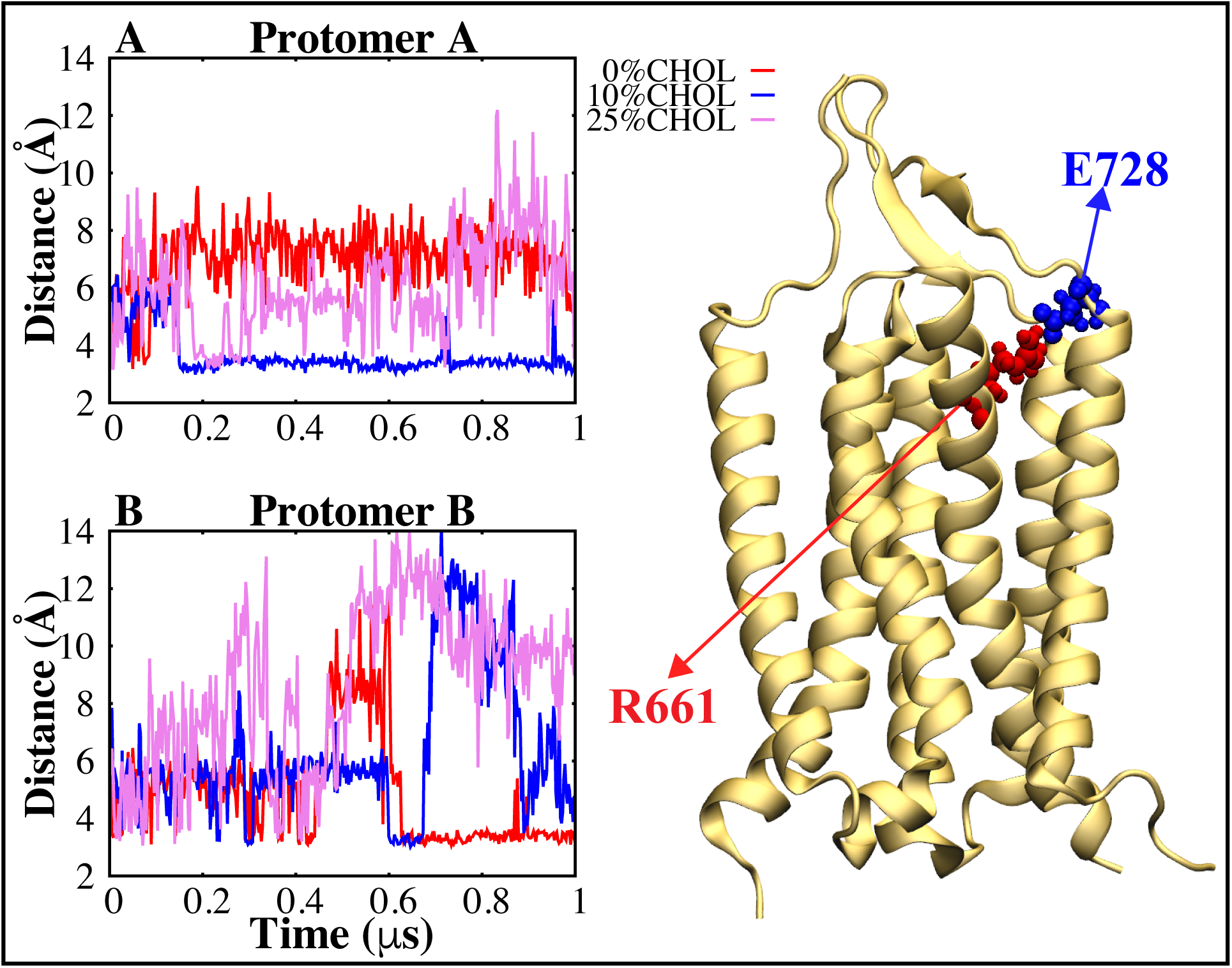
Time series of the salt bridge network between C*α* atoms of R661 and E728 in Protomer A (A) and Protomer B (B). The right panel illustrates a graphical representation of mGluR1, highlighting the salt bridge interactions between R661 (red) and E728 (blue) in the 10% CHOL system for Protomer A.

### Cholesterol in mGluR1 is Localized at the Inter-Protomer Interface and Interacts Preferentially with Transmembrane Helices 1 and 2

Our dynamical network analysis reveals significant differences in correlation patterns between the cholesterol-free system (0% CHOL) and systems containing cholesterol (10% and 25% CHOL) (Fig. 6). Particularly, pronounced differences are observed within the transmembrane helices 1 and 2 (TM1-TM2) of both protomers A and B. The presence of cholesterol induces substantial changes in the correlated motions of residues in these regions, evident by the darker red gradient in Figure 6. This indicates cholesterol’s crucial role in modulating the dynamics and interactions of the transmembrane helices surrounding mGluR1’s interphase. The altered correlations in TM1-TM2 residues upon cholesterol addition suggest cholesterol’s influence on the conformational flexibility and coupling of transmembrane helices, potentially modulating the receptor’s ability to transmit signals across the membrane. ^4^ Remarkably, our analysis reveals that the system with 10% cholesterol (Fig. 6A) exhibits more pronounced differences in residue correlations compared to the 25% cholesterol system (Fig. 6B). This suggests that a relatively low concentration of cholesterol (10%) significantly impacts mGluR1 dynamics, potentially optimizing lipid-protein interactions and stabilizing specific conformational states of the receptor. The preferential impact of cholesterol on the transmembrane helices surrounding mGluR1’s inter-phase highlights this region’s importance in receptor function and regulation. These helices are implicated in dimerization and allosteric communication between mGluR1 protomers.^67,68^ Cholesterol-induced changes in TM1-TM2 dynamics may influence the stability and signaling properties of the mGluR1 dimer, impacting receptor activation and downstream signaling pathways.^67,68^

**Fig. 6.**
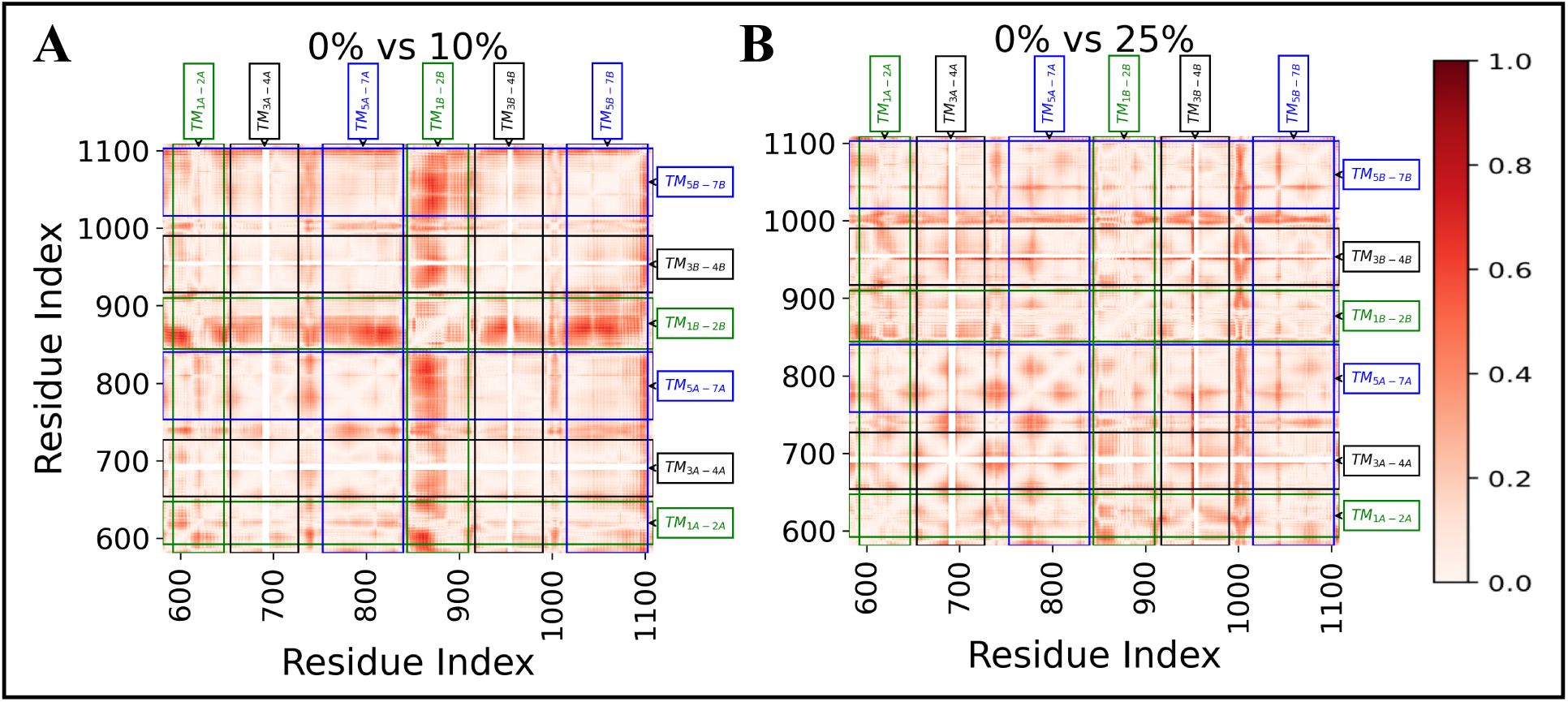
The dynamical network analysis illustrates correlation differences between the 0% cholesterol and 10% cholesterol systems (A), and between the 0% cholesterol and 25% cholesterol systems (B) for simulation set 1. Differences in correlation are shown as a red gradient, where darker shades indicate higher differences, ranging from 0 to 1.

Our findings contribute to the understanding of cholesterol’s critical role in GPCR function and regulation.^4,16,69–71^ Cholesterol interacts with specific regions of GPCRs, such as the cholesterol recognition amino acid consensus (CRAC) motif, modulating their structural and functional properties.^72–75^ The preferential impact of cholesterol on mGluR1’s transmembrane helices aligns with cholesterol-mediated GPCR regulation, emphasizing the significance of considering the lipid environment in investigating GPCR dynamics and function.

In Figure 7, we present a graphical representation highlighting the predominant localization of cholesterol within the interphase of both the 10% and 25% systems. This localization pattern is notably observed between protomers, suggesting a specific affinity for this region (Fig. 7A & B). To further substantiate this finding, our analysis of interhelical angles reveals notable fluctuations, particularly involving TM1 with other helices, as well as TM2 with other helices (Fig. S1, S2). Considering that TM1 and TM2 collectively form the interphase region of the receptor, it’s evident that cholesterol tends to concentrate in this region. This localization within the interphase underscores the significance of cholesterol in influencing the conformational dynamics and structural organization of mGluR1, particularly in regions crucial for receptor function and regulation.

**Fig. 7.**
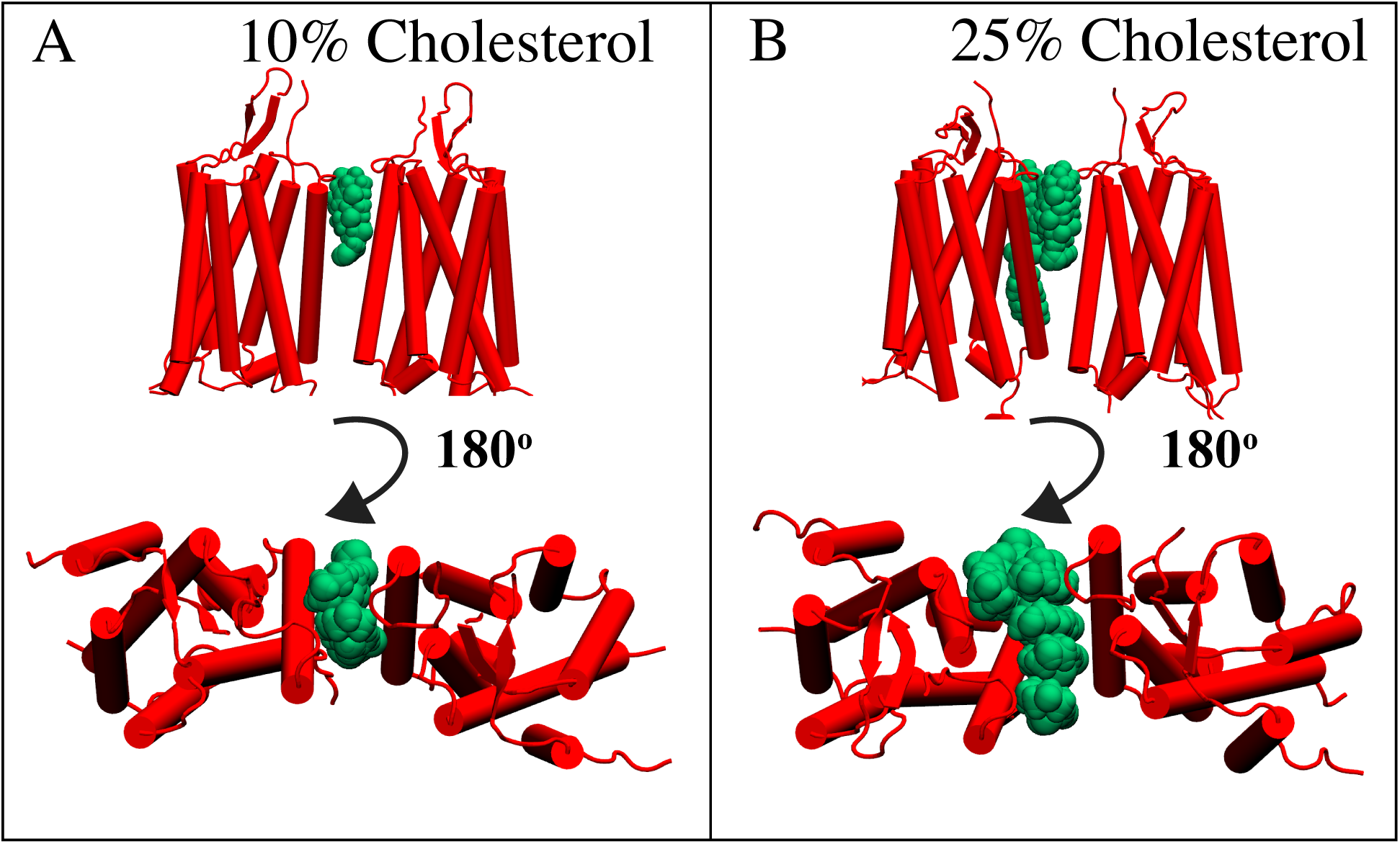
A graphical representation highlighting the preferred localization region of cholesterol (green) within the mGluR1 (red) systems with 10% (A) and 25% (B) cholesterol concentrations.

### Impact of Removing Initial Six Cholesterol Molecules on Receptor Conformational Dynamics

Examining Set 2, which excludes the initial six cholesterol molecules from the crystal structure, offers additional insights. As in Set 1, significant changes in conformational dynamics are most evident when considering the entire mGluR1 protein rather than individual protomers (Fig. 8A-F). Notably, Set 2 reveals a heightened degree of conformational change in the absence of cholesterol (0%), with a similar trend observed for both the 0% and 25% systems, contrasting with the distinct behavior seen in the 10% system, echoing observations from Set 1 (Fig. 8C & F).

**Fig. 8.**
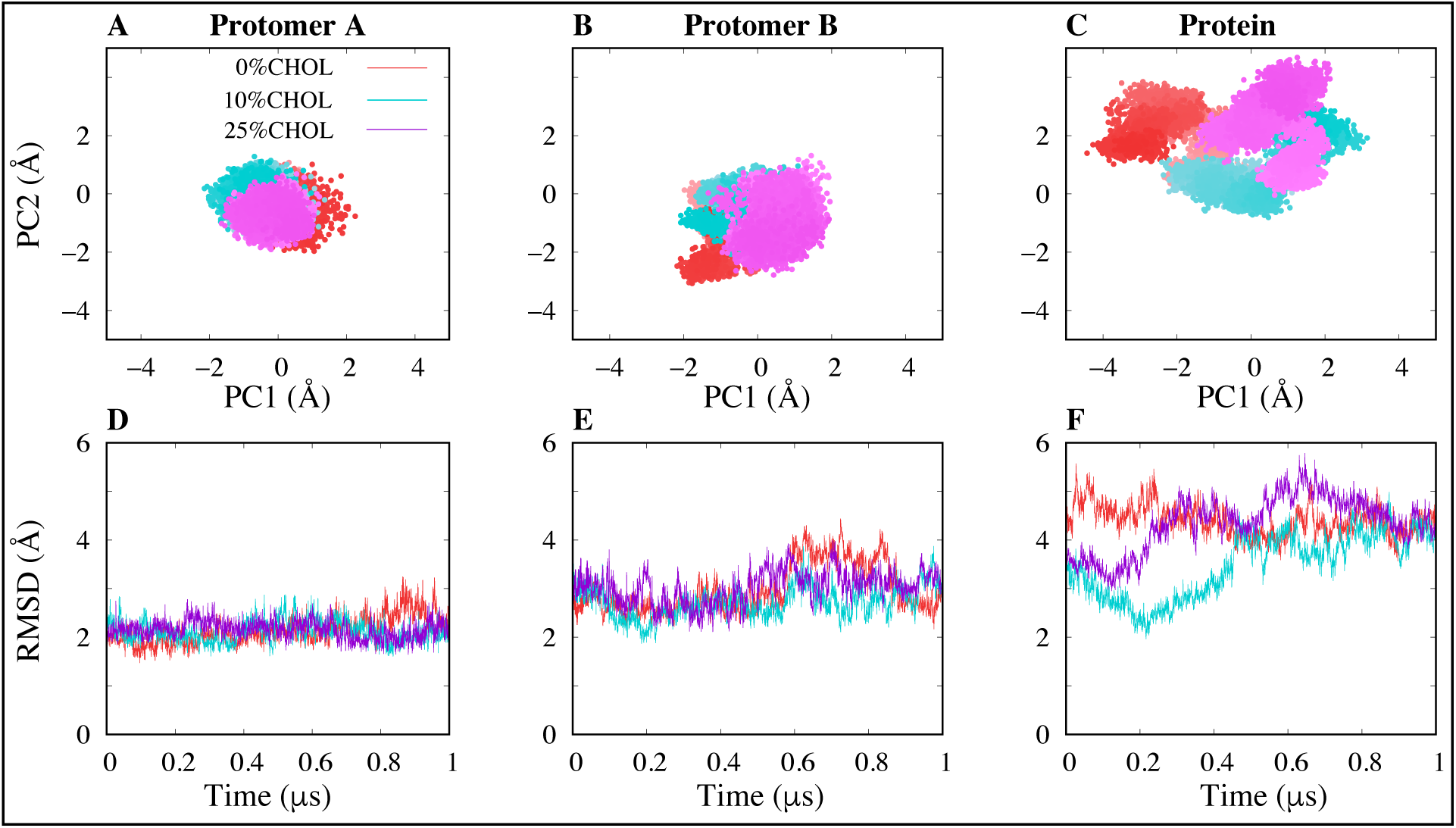
Projections of the principal components (PC’s) 1 and 2 [A-C], and the root mean square deviation analysis [D-F] of mGluR1 in the presence and absence of cholesterol, in the second simulation set.

This indicates that the absence of cholesterol amplifies conformational variability in mGluR1, allowing for increased flexibility within the protein structure. Conversely, the presence of cholesterol, particularly at 10% concentration, influences conformational dynamics differently, suggesting a balance between stabilization and flexibility mediated by cholesterol-lipid interactions.

The RMSD distribution plots support the observations and provide insights into the interplay between individual protomers and the overall behavior of the receptor complex. In Set1, the system with no cholesterol (0% CHOL) displays a distinct high peak in the RMSD distributions of both Protomer A and Protomer B (Fig. 9A & B). However, when considering the whole protein (Fig. 9C), the RMSD distribution shows a lower and broader peak, indicating a more diverse conformational sampling of the receptor complex in the absence of cholesterol. The system with 25% cholesterol (Fig. 9A-C) maintains a consistent mid-range peak in the RMSD distributions across the individual protomers and the whole protein. This suggests that the presence of a high cholesterol concentration induces a stabilization of the receptor’s structure, limiting the extent of conformational fluctuations. In contrast, the 10% cholesterol system (Fig. 9A-C) exhibits a more complex pattern, with the whole protein (Fig. 9C) displaying a lower and broader peak compared to the individual protomers. This observation implies that the intermediate cholesterol concentration may lead to increased conformational diversity in the overall receptor complex, potentially reflecting a more dynamic and adaptable state. The observations from the first simulation set (Set1) highlight the asymmetric behavior of the mGluR1 protomers and the distinct impact of cholesterol on the conformational dynamics of the individual subunits and the receptor as a whole. Interestingly, the patterns observed in Set2 reveal a somewhat different scenario. In this set, the system with no cholesterol (0% CHOL) (Fig. 9D-F) exhibits the lowest peaks in the RMSD distributions of the individual protomers (Protomer A and Protomer B), while the whole protein (Fig. 9F) displays the highest peak. This suggests that in the absence of cholesterol, the receptor complex may adopt a more diverse conformational landscape compared to the individual protomers. Furthermore, the 10% cholesterol system in Set2 (Fig. 9F) shows the lowest peak in the RMSD distribution of the whole protein, revealing a bimodal pattern similar to that observed in the first simulation set. The patterns observed across the two independent simulation sets (Set1 and Set2) underscore the nature of the findings and a relatively consistent reproducibility of the cholesterol-mediated modulation of mGluR1 conformational dynamics. The asymmetric behavior of the protomers and the distinct receptor-level responses to varying cholesterol concentrations highlight the complex and context-dependent nature of the receptor’s structural flexibility.

**Fig. 9.**
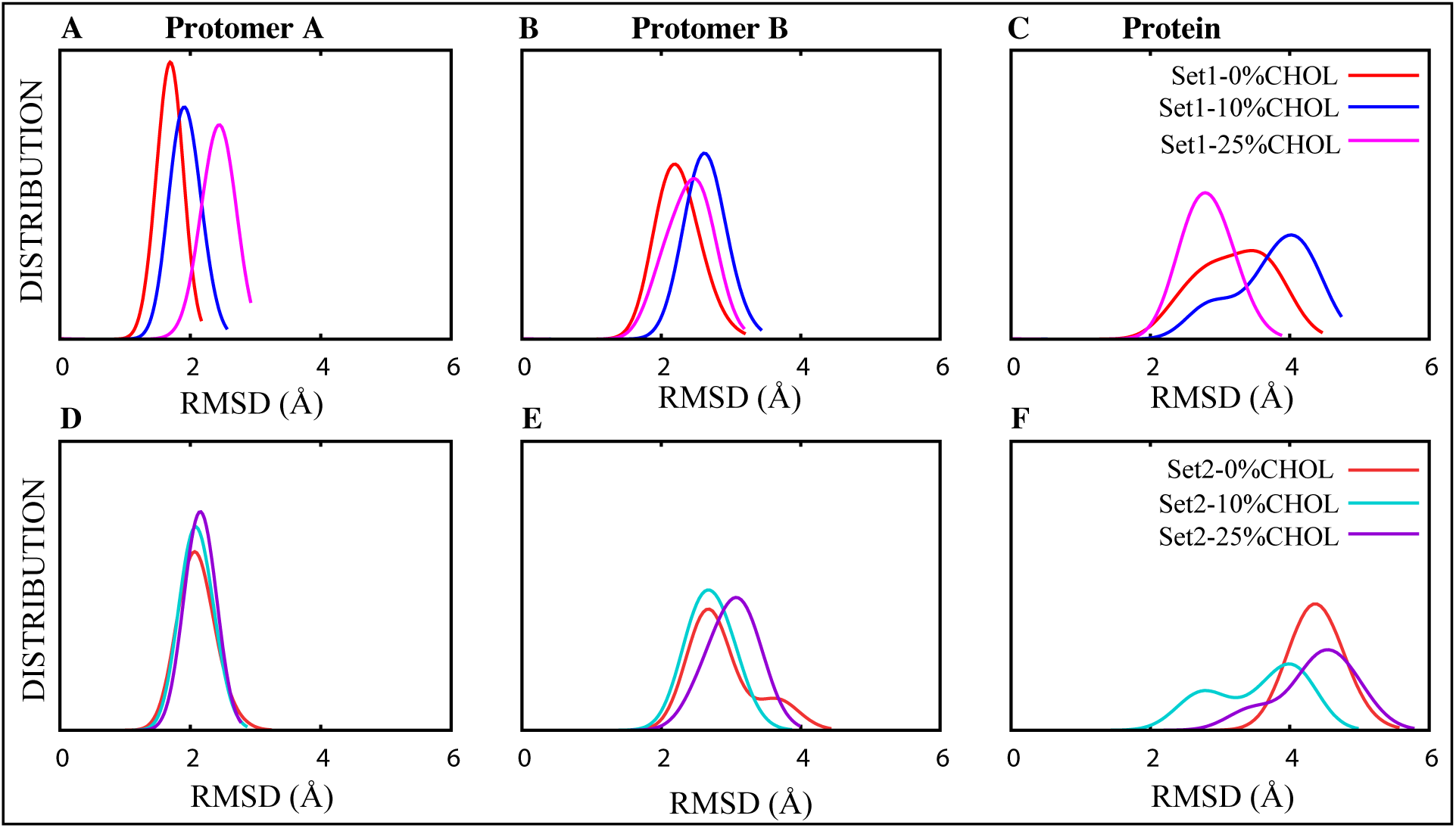
RMSD distribution plots across different cholesterol concentrations in Simulation Set 1 [A-C] and Simulation Set 2 [D-F].

Considering the inter-protomer angle and distance, the observations from the second simulation set (Set2) reveal a somewhat inconsistent yet similar pattern. In this set, the 0% and 25% cholesterol systems exhibit similar behavior, albeit with a higher inter-protomer distance and lower angle (Fig. 10A) compared to Set1 (Fig. 3). Notably, the 10% cholesterol system in Set2 displays a distinct behavior, showing a lower inter-protomer distance and a higher angle, contrasting with Set1 (Fig. 10B). The observed cholesterol-dependent modulation of the inter-protomer distance and angle has important implications for understanding the functional dynamics of mGluR1. The dimeric arrangement of mGluR1 plays a crucial role in its signaling properties and allosteric regulation. Changes in the inter-protomer distance and angle may affect the stability and communication between the mGluR1 protomers, potentially influencing receptor activation and downstream signaling events.

**Fig. 10.**
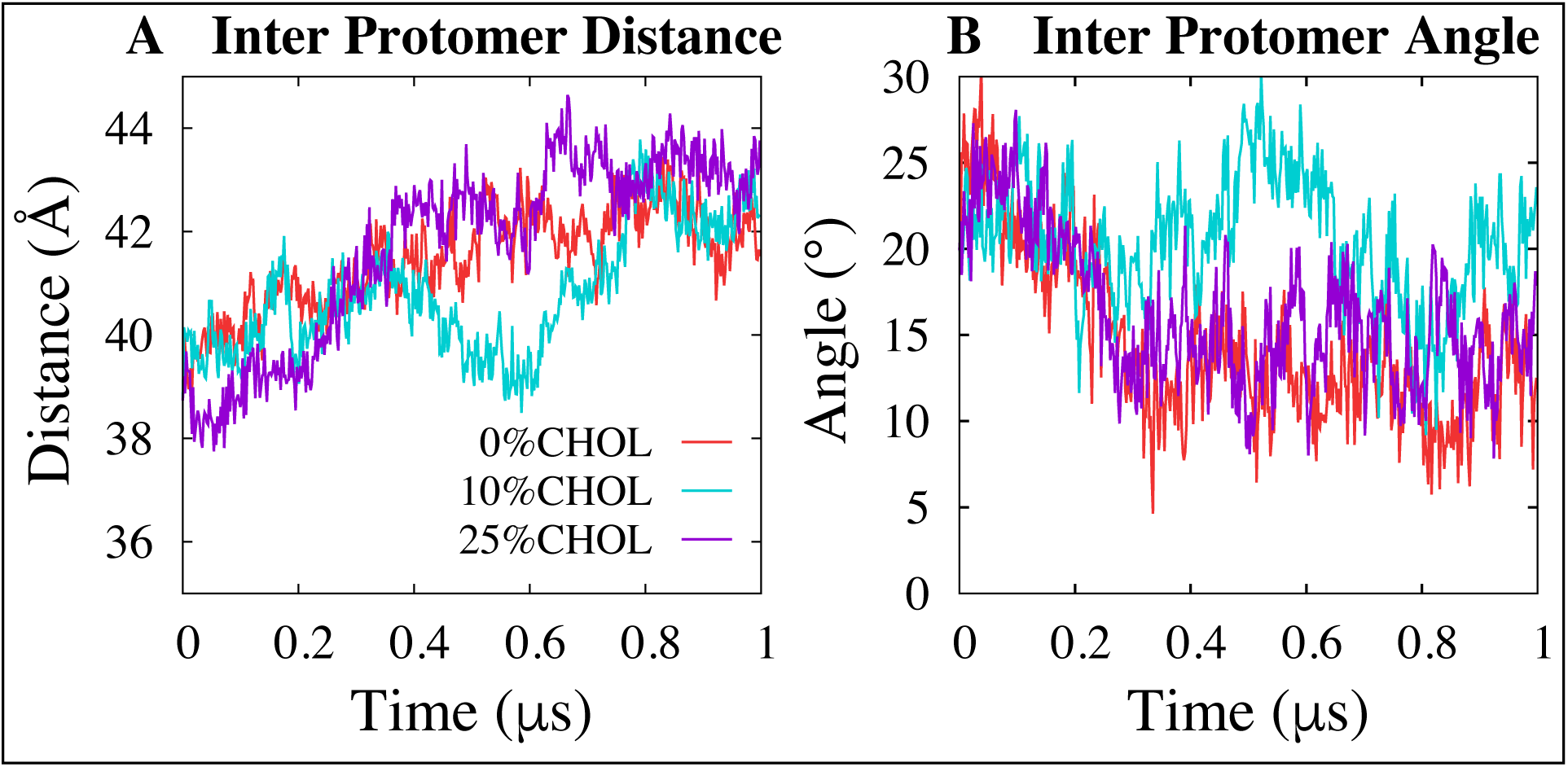
Time series representation of the inter-protomer distance (left panel) and interprotomer angle (right panel) in Simulation Set 2.

Additionally, when analyzing the water density in Set 2, specifically with 10% cholesterol, the presence of water in the lower leaflet persists consistently throughout the entire simulation duration (Fig. 11). In contrast, in the simulations with 0% and 25% cholesterol, there is a noticeable decrease in water density during the last 200 ns compared to the initial 200 ns of the simulations (Fig. 11).

**Fig. 11.**
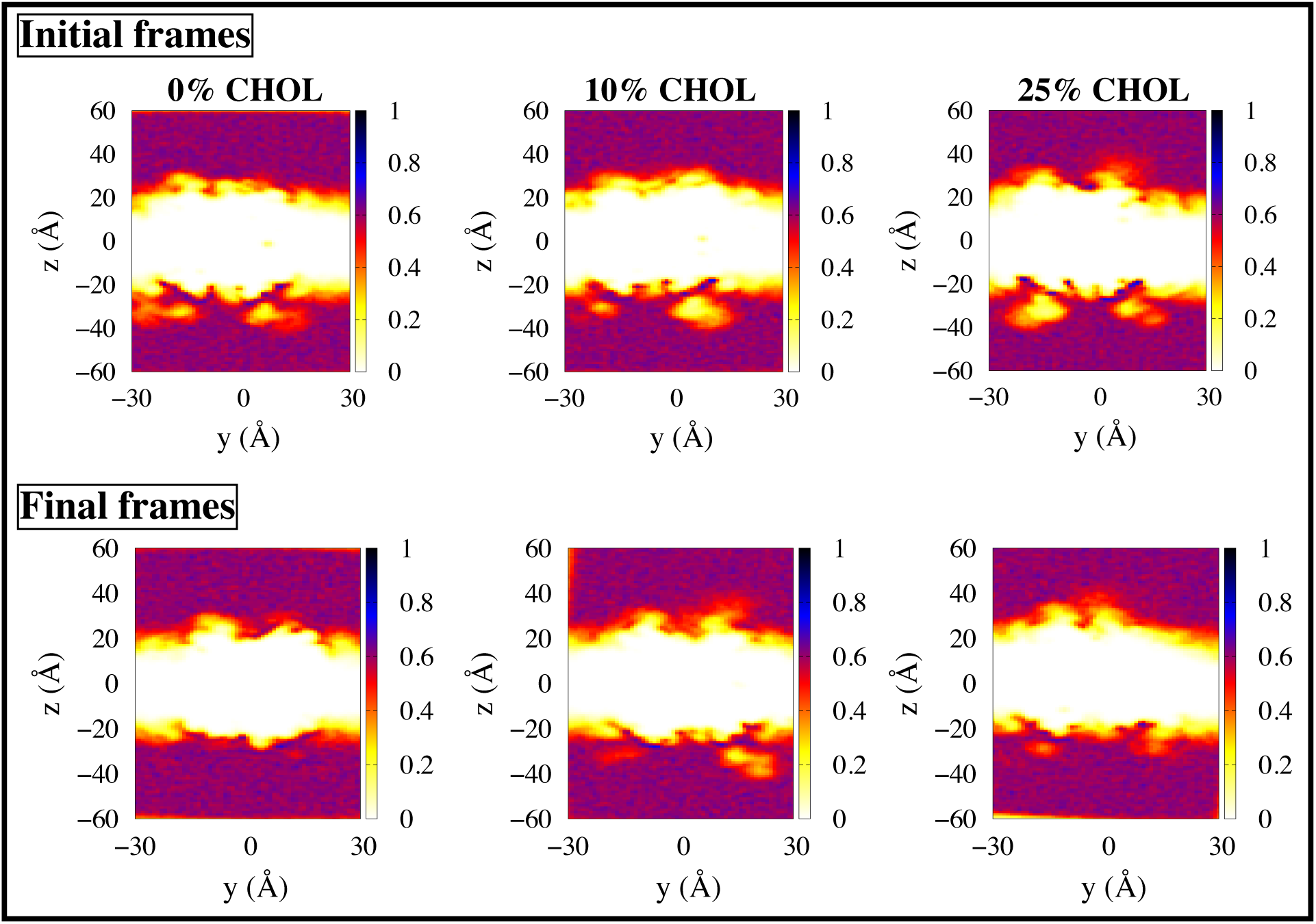
Water density maps illustrate the initial frame (first 200 ns) and final frames (last 200 ns) for Set 1 across 0%, 10%, and 25% cholesterol concentrations. Variations in correlation are represented by a color gradient, ranging from 0 (pale yellow) to 1 (dark blue).

Overall, we note contrasting yet consistent trends across the two distinct simulation sets (Set1 and Set2). While we consistently observe greater conformational changes when considering the entire protein rather than individual protomers, and the 10% system consistently behaving differently, we also observe a reversal in the arrangement. This underscores the significance of the findings and the reproducibility of cholesterol-mediated modulation of mGluR1 conformational dynamics. The asymmetric behavior of the protomers and the distinct responses of the receptor to different cholesterol concentrations underscore the complex and context-dependent nature of the receptor’s structural flexibility. These insights contribute to a deeper understanding of the structure-function relationship of mGluR1 and the pivotal role of cholesterol in regulating its conformational dynamics. This knowledge may have important implications for the development of targeted pharmacological interventions that leverage the cholesterol-mediated modulation of mGluR1 for therapeutic applications.

## Conclusion

In this study, we have investigated the influence of cholesterol on the conformational dynamics of mGluR1. We have observed that cholesterol exerts a more pronounced impact on the overall conformation of mGluR1 compared to individual protomers, indicating a collective effect on the receptor’s structural flexibility. Furthermore, our analysis reveals a preferential localization of cholesterol within the inter-phase of the protein.

Notably, our findings demonstrate that the system with a lower cholesterol concentration (10%) exhibits greater conformational changes compared to those with higher cholesterol levels (25%). This observation suggests a concentration-dependent effect of cholesterol on mGluR1 dynamics, with lower concentrations inducing more pronounced alterations in protein conformation. Moreover, as cholesterol concentration increases, we observe a trend towards a more ordered protein structure, characterized by reduced motion between the helices. This highlights the role of cholesterol in stabilizing specific conformational states of mGluR1, potentially influencing its functional properties.

Additionally, our investigation into the impact of removing the initial six cholesterol molecules from the crystal structure underscores the significance of cholesterol in shaping receptor conformational dynamics. We observe notable alterations in the conformational behavior of mGluR1 upon the removal of these cholesterol molecules, indicating the importance of cholesterol-lipid interactions in maintaining the structural integrity of the receptor.

Overall, our study contributes valuable insights into the role of cholesterol in modulating the conformational dynamics of mGluR1. By elucidating the concentration-dependent effects of cholesterol and its localization within the protein interphase, we enhance our understanding of the structural and functional implications of cholesterol-protein interactions in G protein-coupled receptor signaling. These findings may have broader implications for drug discovery and therapeutic interventions targeting mGluR1 and related receptors in neurological disorders.

## Acknowledgement

The research described in this work received support from various funding sources. Specifically, we acknowledge the National Institute of General Medical Sciences of the National Institutes of Health for their support under award numbers R15GM139140 and R35GM147423. Additionally, this research was supported by the National Science Foundation grant CHE 1945465 and the Arkansas Biosciences Institute. We also appreciate the support provided by the Frontera computing project by LRAC at the Texas Advanced Computing Center (TACC) through Grant CHE21003, made possible by the National Science Foundation award OAC-1818253. Furthermore, we acknowledge the Pittsburgh Supercomputing Center for providing Anton 2 computer time through Grant R01GM116961 from the National Institutes of Health. We extend our gratitude to D.E. Shaw Research for generously making the Anton 2 machine available at PSC. Additionally, this research utilized Stampede by the Extreme Science and Engineering Discovery Environment (allocation MCB150129), which is supported by National Science Foundation grant number ACI-1548562. We also acknowledge the support received from the Arkansas High Performance Computing Center(AHPCC), funded through multiple National Science Foundation grants and the Arkansas Economic Development Commission. Lastly, we appreciate the support provided by the Blue Waters sustained-petascale computing project, which is supported by the National Science Foundation (awards OCI-0725070 and ACI-1238993) and the state of Illinois.

## Supporting Information Available

Figures S1–S3 and Table S1 in Supporting Information provide additional analysis based on our MD simulations, as discussed in the manuscript.

