## Supporting Information for "Cholesterol Dependence of the Conformational Changes in Metabotropic Glutamate Receptor 1"

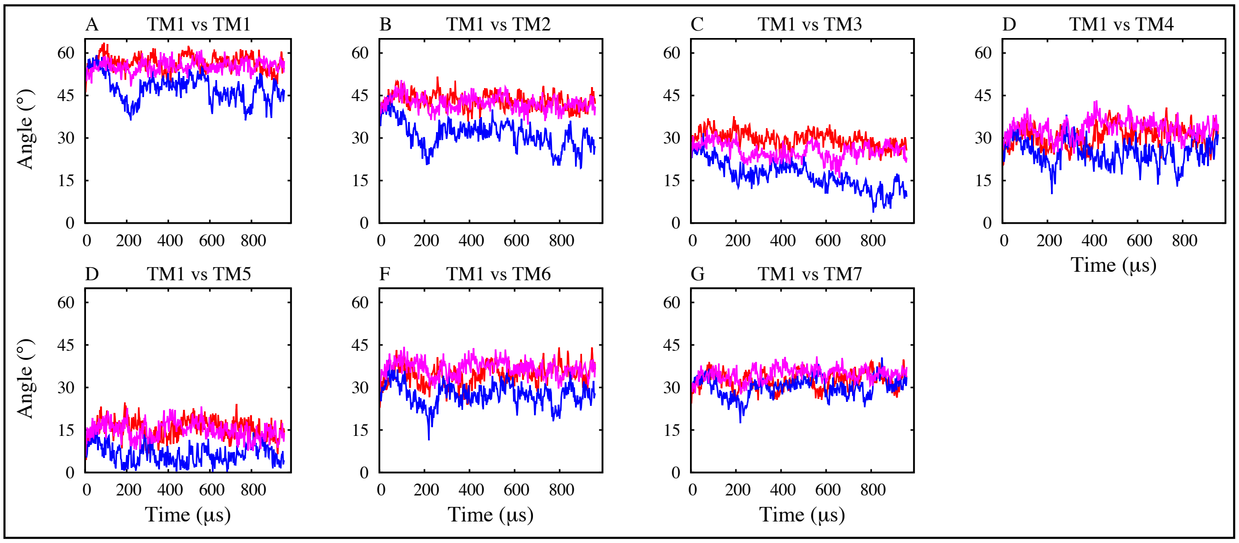


**Fig. S1. Time series plot showing the interhelical angle throughout the simulations for simulation set 1. The interhelical angle between TM1 and TM2-TM7 has been calculated for each frame of the simulation.**

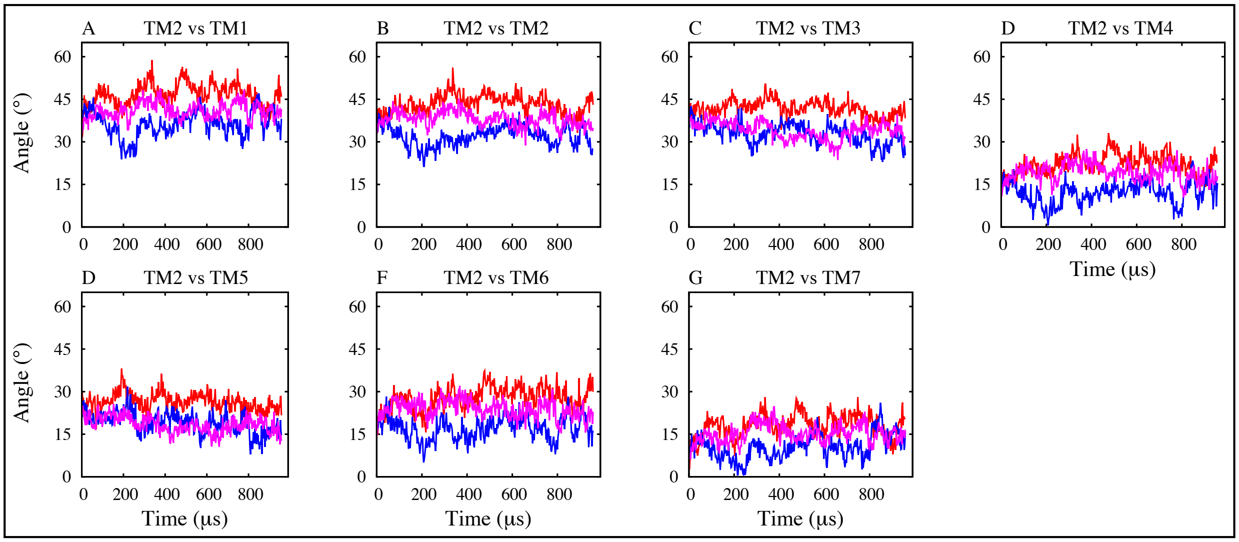


**Fig. S2. Time series plot showing the interhelical angle throughout the simulations for simulation set 1. The interhelical angle between TM2 and TM1, TM3-TM7 has been calculated for each frame of the simulation.**

### Tables

**Table S1: Occupancy (%) of Hydrogen Bond Interactions of All Protomers from Simulation Set 1**

| | **0 % CHOL** | | **10 % CHOL** | | **25 % CHOL** | |
|---|---|---|---|---|---|---|
| System | **A** | **B** | **A** | **B** | **A** | **B** |
| **Y672-T794** | 3 | 12 | 13 | 8 | **98** | 6 |
| **T768-Y792** | 0 | 0 | 0 | 0 | **51** | 0 |
| **S627-K678** | **19** | 59 | **35** | 61 | 68 | 73 |
| **M791-T768** | 94 | 95 | 96 | 95 | **34** | 95 |
| **Y770-N680** | 95 | 96 | 96 | **51** | 98 | **33** |
| **R661-E728** | 3 | 46 | **84** | 13 | 14 | 8 |
| **E783-K834** | 74 | 98 | 97 | 97 | 94 | 73 |
